# Selective sweep and phylogenetic models for the emergence and spread of pyrimethamine resistance mutations in *Plasmodium vivax*

**DOI:** 10.1101/433128

**Authors:** Ayaz Shaukat, Qasim Ali, Timothy Connelley, Muhammad Azmat Ullah Khan, Mushtaq A. Saleem, Imran Rashid, Neil D. Sargison, Umer Chaudhry

**Affiliations:** Faculty of Life Sciences, University of Central Punjab, Lahore, Pakistan; Department of Parasitology, University of Veterinary and Animal Sciences Lahore, Pakistan; University of Edinburgh, The Roslin Institute, Easter Bush Veterinary Centre, Roslin, Midlothian, EH25 9RG, UK

**Keywords:** Pyrimethamine resistance, Plasmodium vivax, Selective sweeps

## Abstract

Pyrimethamine resistance is a major concern for the control of human haemoprotozoa, especially *Plasmodium* species. Currently, there is little understanding of how pyrimethamine resistance developed in *Plasmodium vivax* in the natural field conditions. Here, we present first time the evidence of positive selection pressure on a dihydrofolate reductase locus and its consequences on the emergence and the spread of pyrimethamine resistance in *P. vivax* in the Punjab province of Pakistan. First, we examined the pyrimethamine resistance locus in 38 *P. vivax* populations to look for evidence of positive selection pressure in human patients. The S58R (AG<u>A</u>)/S117N (A<u>A</u>C) double mutation was most common, being detected in 10/38 populations. Single mutation S117N (A<u>A</u>C), I173L (<u>C</u>TT) and S58R (AG<u>A</u>) SNPs were detected in 8/38, 2/38 and 1/38 populations, respectively. The F57L/I (TT<u>A</u>/<u>A</u>T<u>A</u>) and T61M (A<u>T</u>G) SNPs were not detected in any population examined. Although both soft and hard selective sweeps have occurred with striking differences between populations, there was a predominance of hard sweeps. A single resistance haplotype was present at high frequency in 9/14 populations, providing a strong evidence for the single emergence of these mutations. In contrast, 5/14 populations carried multiple resistance haplotypes at high frequencies, providing an evidence of the emergence of resistance by recurrent mutations, characteristics of soft selective sweeps. Our phylogenetic relationship analysis suggests that S58R (AG<u>A</u>)/S117N (A<u>A</u>C) and S117N (A<u>A</u>C) mutations arose multiple times from a single origin and spread to multiple different cities in the Punjab province through gene flow. Interestingly, the I173L (<u>C</u>TT) mutation was present on a single haplotype, suggesting that it arises rarely and has not spread between cities. Our work shows the need for responsible use of exiting and new antimicrobial drugs and their combinations, control the movement of infected patients and mosquito vector control strategies.

## 1. Introduction

Malaria is a significant source of morbidity and mortality especially in pregnant women and young children (Petersen et al., 2011). Despite causing probably 72 to 80 million-malaria cases annually, *P. vivax* usually produces less severe symptoms and has not received as much attention as *P. falciparum*. Nevertheless, *P. vivax* leads to a disabling disease that can be fatal and exacts a similar economic burden to *P. falciparum*. Furthermore, the severity of disease caused by *P. vivax* is increasing in south Asia (Conway, 2007).

Practically all knowledge of the genetics of pyrimethamine drug resistance has been based on candidate gene studies of dihydrofolate reductase. The *Plasmodium vivax* dihydrofolate reductase locus has been analysed in susceptible and resistant isolates and single nucleotide polymorphisms (SNPs) have been observed at codons F57L/I (TT<u>A</u>/<u>A</u>T<u>A</u>), S58R (AG<u>A</u>), T61M (A<u>T</u>G), S117N/T (A<u>A</u>C/A<u>C</u>C) and I173L/F (<u>C</u>TT/<u>T</u>TT) (de Pecoulas et al., 1998; Lee et al., 2010). This provides a strong evidence that these mutations in the dihydrofolate reductase locus are important determinants of pyrimethamine resistance in *P. vivax*. Further functional analysis has demonstrated that *P vivax* dihydrofolate reductase resistant strains do not allow the uptake of pyrimethamine in *Saccharomyces cerevisiae* and *Escherichia coli* models, confirming that dihydrofolate reductase confers resistance to the pyrimethamine drug class (Hastings et al., 2005; Hastings et al., 2004; Hastings and Sibley, 2002). The pyrimethamine resistance SNPs have been further investigated in different geographical regions and evidence exists that the amino acid substitutions in codons F57L/I (TT<u>A</u>/<u>A</u>T<u>A</u>), S58R (AG<u>A</u>), T61M (A<u>T</u>G), S117N/T (A<u>A</u>C/A<u>C</u>C) and I173L/F (<u>C</u>TT/<u>T</u>TT) can be responsible for resistance to this drug class (Auliff et al., 2006; Huang et al., 2011).

Although, some initial progress has been made in elucidating the molecular genetics of pyrimethamine resistance in *P. vivax*, there are very few published reports investigating the populations genetics of the pyrimethamine resistance mutations (Alam et al., 2007; Hawkins et al., 2008), and it is still unclear how resistance develops and spreads in the field. Hypothetically the development of resistance depends on multiple factors including: (a) the intensity of positive selection pressures on the candidate locus due to treatment compliance, or fitness costs associated with the resistance mutations; and (b) emergence of resistance alleles due to high rates of mutations followed by the spread by gene flow (Chaudhry, 2015; Petersen et al., 2011).

Selective sweep models could potentially answer how antimicrobial resistance mutations emerge in parasite populations; whereby use of antimicrobial drugs provides a positive selection pressure for adaptive mutations in the resistance candidate loci of the parasite populations, resulting in selective sweeps at the loci under selection. There are two different types of selective sweep models. A ‘hard’ selective sweep is characterised by a single resistance haplotype rising at high frequency in each parasite population from a single mutation. This allows little time for recombination to break up the initial haplotype on which the resistance mutation appeared, therefore, the genetic footprint of selection is expected to involve a reduction in polymorphism around the locus after selection. A ‘soft’ selective sweep is characterised by the presence of multiple resistance haplotypes at high frequency in each population derived from either recurrent mutation appearing after the onset of selection, or pre-existing mutations before the onset of selection. The genetic footprint of selection would not be expected to include a marked reduction in polymorphism around the locus under selection (Chaudhry, 2015).

Understanding of adaptive mutations in response to selection can help to show their spread. There are two phylogenetic models in which adaptive mutations can spread in the populations that are under selection. A new resistance mutation could arise from a single origin, become fixed by selection and then spread through parasite populations by migration, likely a consequence of the flow of drug resistance alleles. Alternatively, resistance mutations could repeatedly arise from multiple origins, become fixed by selection and migrate between parasite populations as a result of the flow of drug resistance alleles (Chaudhry, 2015).

Previously most targeted studies characterising pyrimethamine resistance have been undertaken in *Plasmodium falciparum* (Petersen et al., 2011). There is currently little understanding of how pyrimethamine resistance developed, emerged and spread in *P. vivax*. In the present study, we examined hard and soft selective sweep models in which single resistance mutations or multiple independent recurrent resistance mutations could emerge in *P. vivax* and then spread by migration. This helps to explain why pyrimethamine resistance is so common in *Plasmodium* parasites and suggests that the emergence of pyrimethamine resistance is likely when large size parasite populations are exposed to antimicrobial drugs.

## 2. Materials and Methods

### 2.1 Parasite material and morphological identification of Plasmodium at genus level

Malaria patients referred to the Chughtai diagnostic laboratory in the Punjab province were invited to participate in this study. The procedures involved in sample collection by venipuncture are minimal. Discussions have been held with key administrative and community leaders to raise awareness of the study. Trained paramed workers under the supervision of local collaborator and doctor have taken samples. This rigorous approach has been continued throughout the study. Blood samples have been collected during peak malaria transmission seasons between April and October 2016 and 2017. The study included patients of all age groups with malaria symptoms including vomiting, fever, headache, chills, sweats, nausea and fatigue. Institutional Review Board of the University of Central Punjab, Pakistan approved the study. 5 ml of intravenous blood was drawn into an EDTA tube by venipuncture from patients giving written consent and stored at -20 □C. 3% Giemsa stained thin and thick blood smears were prepared and examined by a trained technician under the oil immersion (1000 x) for the diagnosis of *Plasmodium*, according to WHO guidelines (Asif, 2008).

### 2.3 Genomic DNA isolation and molecular identification of Plasmodium species

About 50 μL of each blood sample from 38 malaria positive patients was considered to be individual population. The gDNA was extracted according to the protocols described in the TIANamp blood DNA kit (Beijing Biopeony Co. Ltd). For DNA preparations, 1 μL from each gDNA preparation was taken in 4 μL of ddH_2_O to make a 1:5 dilution for use as PCR template. Dilutions of aliquots of only ddH_2_O were made in parallel for use as negative controls. For the species confirmation of *P. vivax*, deep amplicon sequencing (Illumina MiSeq) of the 18s rDNA was performed (Shaukat et al., 2018).

### 2.4 Deep amplicon sequencing of the dihydrofolate reductase locus

A 468bp fragments encompassing parts of F57L/I (TT<u>A</u>/<u>A</u>T<u>A</u>), S58R (AG<u>A</u>), T61M (A<u>T</u>G), S117N/T (A<u>A</u>C/A<u>C</u>C) and I173L/F (<u>C</u>TT/<u>T</u>TT) SNPs of 38 *P. vivax* positive populations were used for deep amplicon sequencing of the dihydrofolate reductase locus. Primers has been modified from the normal primers set (PvdhfrS _For, PvdhfrS_Rev) previously described by Auliff et al. (2006) and Brega et al. (2004). Adapters were added to each primer to allow the successive annealing and N is the number of random nucleotides included between the primers and adopter sequence (Supplementary Table S1).

Four forward (PvdhfrS_For, PvdhfrS_For-1N, PvdhfrS_For-2N, PvdhfrS_For-3N) and four reverse primers (PvdhfrS_Rev, PvdhfrS_Rev-1N, PvdhfrS_Rev-2N, PvdhfrS_Rev-3N) were mixed in equal proportion. The primers were then used for the first round PCR under the following conditions: 5X KAPA HiFi Hot START Fidelity buffer, 10 mM dDNTs, 10 uM forward and reverse adopter primer, 0.5 U KAPA HiFi Hot START Fidelity Polymerase (KAPA Biosystems, USA), 13.25 μL ddH_2_O and 1 μL of lysate. The thermocycling condition of the PCR were 95°C for 2 minutes, followed by 35 cycles of 98°C for 20 seconds, 60°C for 15 seconds, 72°C for 15 seconds and a final extension of 72°C for 5 minutes. PCR products were purified with AMPure XP Magnetic Beads (1X) using a special magnetic stand and plate according to the protocols described by Beckman coulter, Inc.

After the purification, a second round PCR was performed using eight forward and twelve reverse barcoded primers. Repetitions of the same forward and reverse barcoded primers in different samples were avoided. The second round PCR conditions were: 5X KAPA HiFi Hot START Fidelity buffer, 10 mM dDNTs, 0.5 U KAPA HiFi Hot START Fidelity Polymerase (KAPA Biosystems,USA), 13.25 μL ddH_2_O and 2 μL of first round PCR product as DNA template. The barcoded forward (N501 to N508) and reverse (N701 to N712) primers (10 uM each) were obtained from Illumina MiSeq protocols (Supplementary Table S2). The thermocycling conditions of the PCR were 98°C for 45 seconds, followed by 7 cycles of 98°C for 20 seconds, 63°C for 20 seconds, and 72°C for 2 minutes. PCR products were purified with AMPure XP Magnetic Beads (1X) according to the protocols described by Beckman coulter, Inc. The pool library was checked with a KAPA qPCR library quantification kit (KAPA Biosystems, USA). The prepared library was then run on an Illumina MiSeq Sequencer using a 500-cycle pair end reagent kit (MiSeq Reagent Kits v2, MS-103-2003) at a concentration of 15 nM with addition 25% Phix Control v3 (Illumina, FC-11-2003). The MiSeq separated all sequences by sample during post-run processing using recognised indices and to generate FASTAQ files.

### 2.5 Deep amplicon sequencing data handling and bioinformatics filter to remove sequencing induced mutations

MiSeq data were analysed with our own adapted pipeline. Briefly, NCBI BLASTN search was used to generate consensus sequences of the dihydrofolate reductase locus. Consensus sequences were built from FASTA files using Geneious Pro 5.4 software (Drummond AJ, 2012). Those MiSeq sequences that did not hit with the dihydrofolate reductase consensus sequences were discarded as contaminants or artifacts. Overall millions of pyrimethamine susceptible and resistance reads of the dihydrofolate reductase locus were generated from the MiSeq data set of the 38 *P. vivax* populations (Table 1). Polymorphisms appearing more than once in the data set were expected to be real, whereas polymorphisms that only occurred once were considered as artefacts due to sequencing errors and were removed as previously described by Chaudhry et al. (2015) and Redman et al. (2015).

**Table 1.**
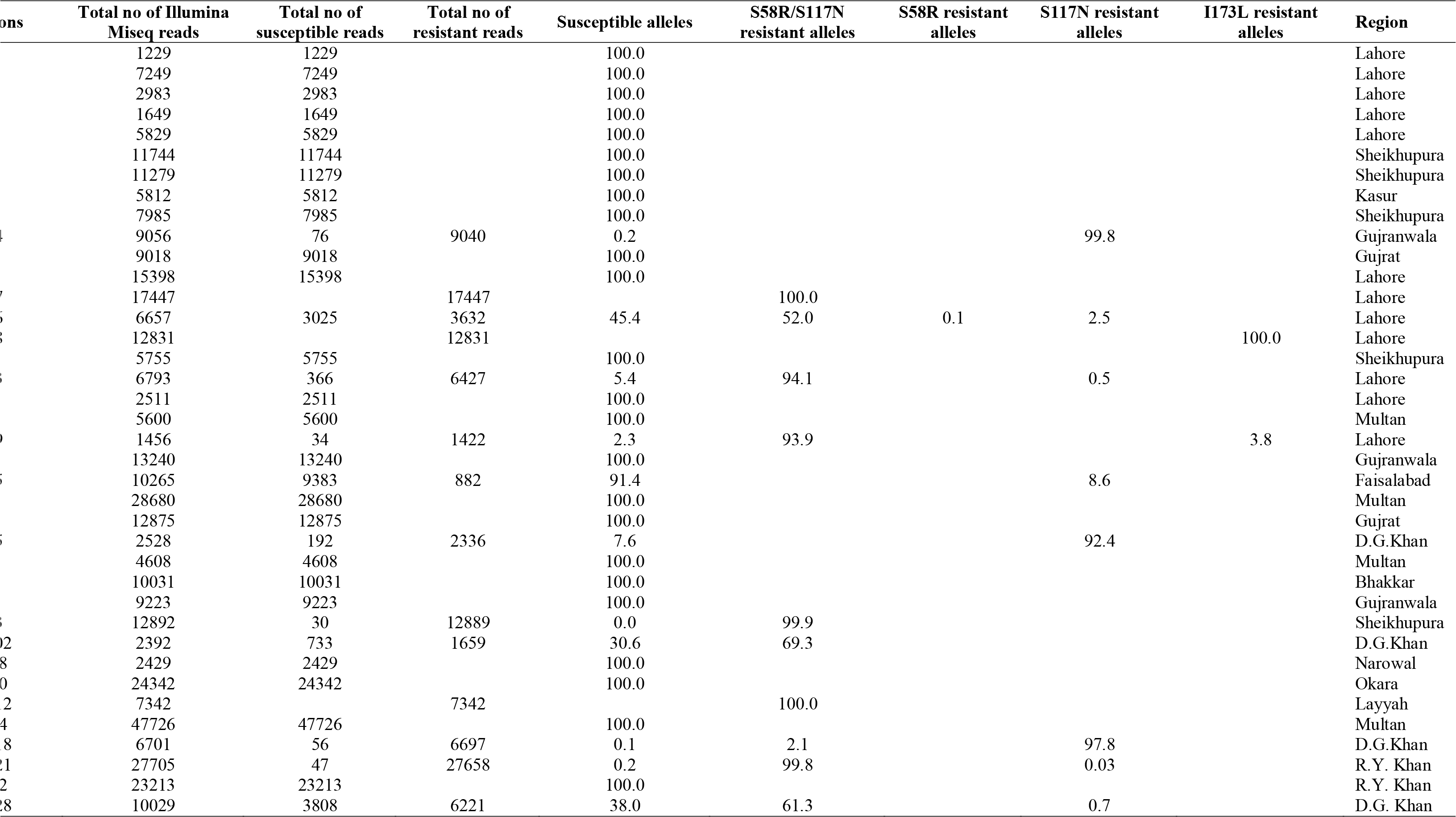
Relative allele frequencies of the dihydrofolate reductase pyrimethamine resistance associated mutations in thirty-eight *P. vivax* populations from the province of Pakistan. Each *P. vivax* population was characterized by the susceptible [S58R(AG<u>C</u>), S117N(A<u>G</u>C), I173L (A<u>T</u>T)] and resistance mutations AG<u>A</u>)/S117N(A<u>A</u>C), S58R(AG<u>A</u>), S117N(A<u>A</u>C) and I173L(<u>C</u>TT)]. The relative allele frequency of the resistant versus susceptible SNPs are based on the entification using Illumina MiSeq deep amplicon sequencing technology.

### 2.6 Genetic diversity estimation and selective neutrality test of the dihydrofolate reductase locus

Genetic diversity estimation and selective neutrality tests were calculated using the DnaSP 5.10 software package (Librado and Rozas, 2009). Briefly, sequence polymorphism estimation is calculated through the haplotype frequency (H_f_), haplotype diversity (H_d_), nucleotide diversity (π), the mean number of pairwise differences (k), the number of segregating sites (S) and the mutation parameter based on an infinite site equilibrium model and the number of segregating sites (θ_S_). Tests for selective neutrality were analysed to determine whether the observed frequency distribution of sequence polymorphism departed from neutral expectations. The neutrality tests included Tajima’s D (Tajima, 1989) and Fay and Wu’s H (Fay and Wu, 2000) methods. Confidence limits and p-values were obtained by coalescent simulations of 10,000 replicates.

### 2.7 Split and the network tree analysis of pyrimethamine resistance associated SNPs in the dihydrofolate reductase locus

A split tree of dihydrofolate reductase haplotypes was generated based on the genetic distance model (JukesCantor) using the UPGMA method employed in SplitTrees4 software (Huson and Bryant, 2006). A split tree of the dihydrofolate reductase haplotypes was also constructed using UPGMA method in MEGA5 software (Tamura, 2011). The program jModeltest 12.2.0 was used to select the appropriate model of nucleotide substitutions for UPGMA analysis (Posada, 2008). According to Bayesian information criterion, the best scoring was JukesCantor (JK+G+I). Branch supports were obtained using 1000 bootstraps of the data. The most probable ancestral node was determined by rooting the tree to a closely related outgroup (*Plasmodium falciparum*). The network tree of dihydrofolate reductase haplotypes was produced based on the Neighbour Joining algorithm built on a sparse network and the epsilon parameter is set to zero default in Network 4.6.1 software (Fluxus Technology Ltd.).

## 3. Results

### 3.1 Validation of deep amplicon sequencing using known proportions of resistant and susceptible dihydrofolate reductase alleles

Before applying the deep amplicon sequencing method to large scale surveys, it is necessary to show that the resistance allele frequency generated by Illumina MiSeq accurately reflects the genotypic frequency. The dihydrofolate reductase susceptible and resistance isolates were selected and run individually through Illumina MiSeq to identify the homozygous susceptible alleles [S: S58R(AG<u>C</u>), S117N(A<u>G</u>C), I173L (A<u>T</u>T)] and homozygous resistant alleles [R: S58R (AG<u>A</u>), S117N (A<u>A</u>C) and I173L (<u>C</u>TT)]. Once identified, pools of gDNA were made based on the known allele frequencies of susceptible and resistance mutations. These were prepared for deep amplicon sequencing and analysis as described above (section 2.4). There was no statistically significant difference between the expected and observed frequencies of susceptible and resistance alleles in a chi-square test, which suggests that the method was accurate in measuring resistance allele frequencies (Fig. 1). For the pools of 100% susceptible or resistance alleles (Mix1:S-100, Mix2: R(S117N)-100, Mix3: R(I173N)-100], the expected and observed results were perfectly matched (Fig. 1). For pools of 50% or 25% susceptible and 50% or 75% resistance alleles [Mix4:S+R(S117N)-50/50, Mix5:S+ R(I173N)-50/50 or Mix6:S+R(S58R/S117N-25/75], the expected and observed results were nearly accurate, with some variations between replicates (Fig. 2). Nevertheless, the data provided a sufficient estimate of allele frequencies appropriate for the work undertaken in this study.

**Fig. 1.**
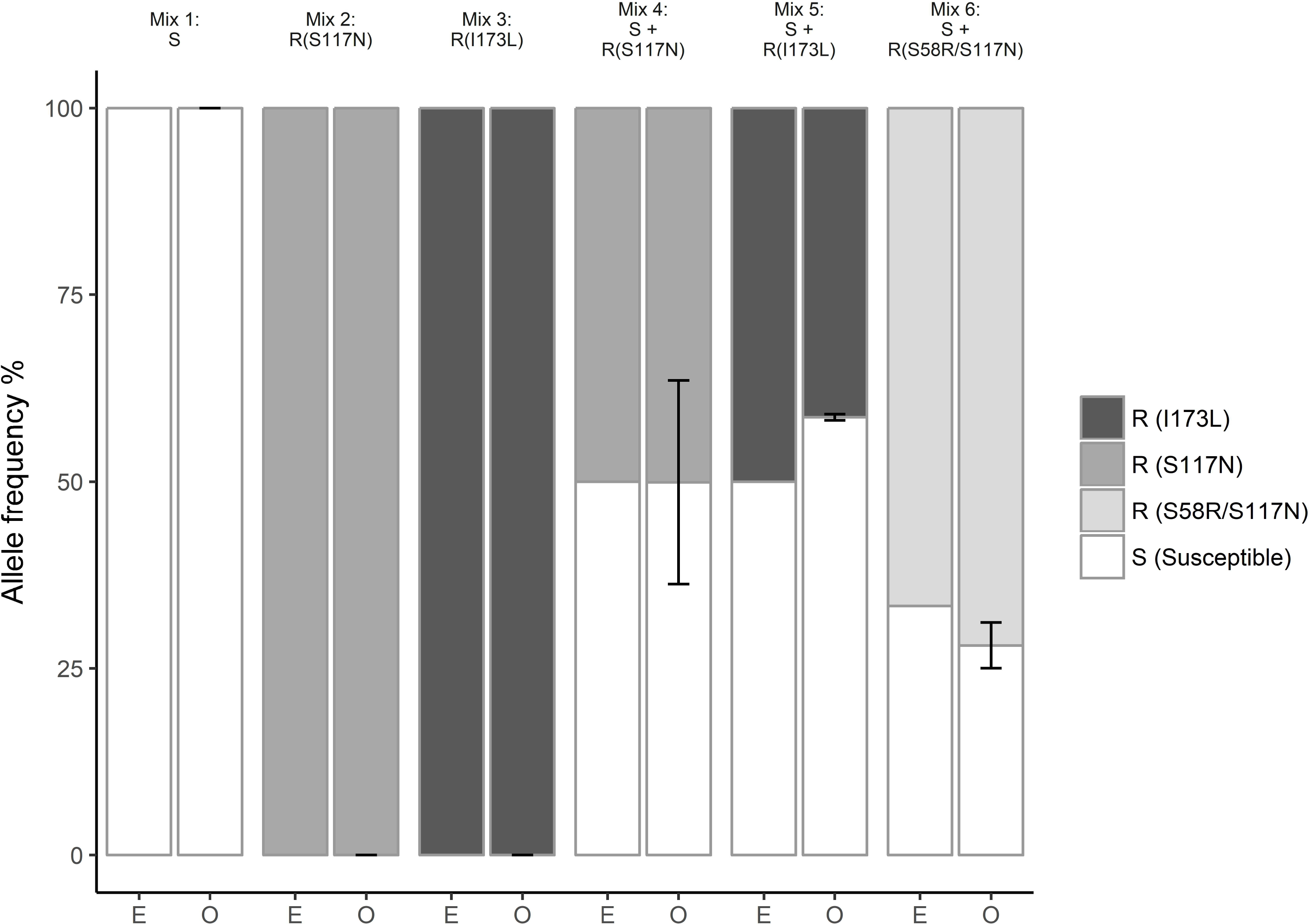
Average frequency of pyrimethamine resistance in mock pools was made from different mixing of the homozygous susceptible alleles [S: S58R(AG<u>C</u>), S117N(A<u>G</u>C), I173L (A<u>T</u>T)] and homozygous resistant alleles [R: S58R (AG<u>A</u>), S117N (A<u>A</u>C), I173L (<u>C</u>TT)]. The symbol mix represents the mixing of homozygous susceptible-S and homozygous resistant-R alleles. In the X-axis, E represents the expected allele frequencies and O represent the observed allele frequencies. White shade indicates susceptible alleles [S: S58R(AG<u>C</u>), S117N(A<u>G</u>C), I173L (A<u>T</u>T)], light grey shade indicates S58R (AG<u>A</u>)/S117N (A<u>A</u>C) resistance alleles, medium grey shade indicates S117N (A<u>A</u>C) resistance allele and dark black shade indicate I173L (<u>C</u>TT) resistance allele. Error bars for observed frequencies represent the standard error of the mean. The mixing of the homozygous susceptible and homozygous resistant were not statistical significant (Chi-square test; Mix1: exact match; Mix2: exact match; Mix3: exact match; Mix4: χ2(1)=1.0089×10-30, p=1; Mix5: χ2(1)=1.1719, p=0.28; Mix6: χ2(1)=0.4259, p=0.51).

**Fig. 2.**
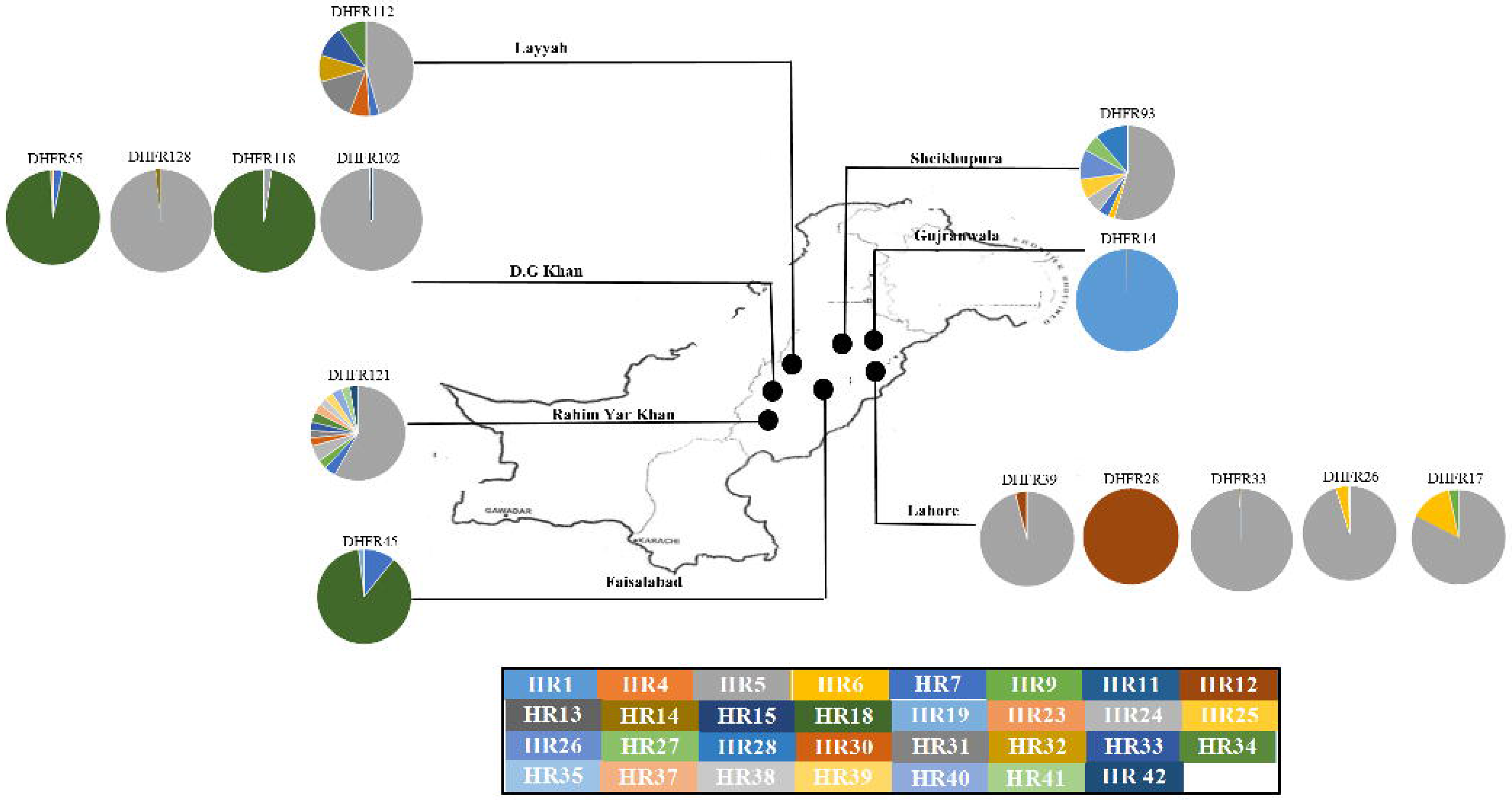
Relative allele frequencies of thirty-one individual resistant haplotypes in fourteen Populations. Susceptible haplotypes were not indicated in the figure but shown in Table 1. The colors in the pie chart circles indicate the haplotype frequency and their distribution in each individual population in the insert map.

### 3.2 Occurrence of pyrimethamine resistance associated SNPs in P. vivax populations

A 468bp fragment of the *P. vivax* dihydrofolate reductase locus was amplified from 38 populations across the Punjab province of Pakistan (Table 1). The relative resistance allele frequencies of five mutations [F57L/I (TT<u>A</u>/<u>A</u>T<u>A</u>), S58R (AG<u>A</u>), T61M (A<u>T</u>G), S117N/T (A<u>A</u>C/A<u>C</u>C) and I173L/F (<u>C</u>TT/<u>T</u>TT)were determined by deep amplicon sequencing. The data set that was generated comprised mostly of three resistance associated SNPs [S58R (AG<u>A</u>), S117N (A<u>A</u>C) and I173L (<u>C</u>TT)] (Table 1). The double mutations of S58R (AG<u>A</u>) and S117N (A<u>A</u>C) were identified in ten populations at frequencies between 2.1 and 100%. The S117N (A<u>A</u>C) SNP was identified in eight populations at frequencies between 0.03 and 99.8%. The I173L (<u>C</u>TT) SNP was detected in two populations at frequencies between 3.8- 100% (Table 1). The pyrimethamine resistance associated SNP S58R (AG<u>A</u>) was present in one population at very low frequency of 0.1% and F57L/I (TT<u>A</u>/<u>A</u>T<u>A</u>) or T61M (A<u>T</u>G) SNPs were not detected in any population examined. Overall results indicated the evidence of positive selection pressure on a dihydrofolate reductase locus.

### 3.3 Dihydrofolate reductase locus diversity and genetic footprint of selection in P. vivax populations

The genetic diversity and neutrality analysis of the dihydrofolate reductase locus was assessed from the 14 selected populations encompassing the S58R (AG<u>C</u>/AG<u>A</u>), S117N (A<u>G</u>C/A<u>A</u>C), I173L (A<u>T</u>T/<u>C</u>TT) resistance mutations (Table 1). Based on the analysis of each population separately, *P. vivax* showed a high level of genetic diversity with a high haplotype diversity (**H_d_**) range from 0.159 to 0.822, a high number of polymorphic sites (S) at all loci, and with a number of alleles per locus ranging from 1 to 33 (Table 2). Overall, there was little difference in the genetic diversity between 14 populations except DHFR134. There was a significant departure of Fay and Wu 's (*H*) and Tajima (*D*) statistic from neutrality in five populations (DHFR17, DHFR45, DHFR93, DHFR112, DHFR121) and a single haplotype was at fixation in the DHFR28 population (Table 2). Five populations had a high frequency of dihydrofolate reductase resistance-conferring haplotypes (Fig. 1), providing evidence of the genetic footprint of selection at the *P. vivax* dihydrofolate reductase locus.

**Table 2:**
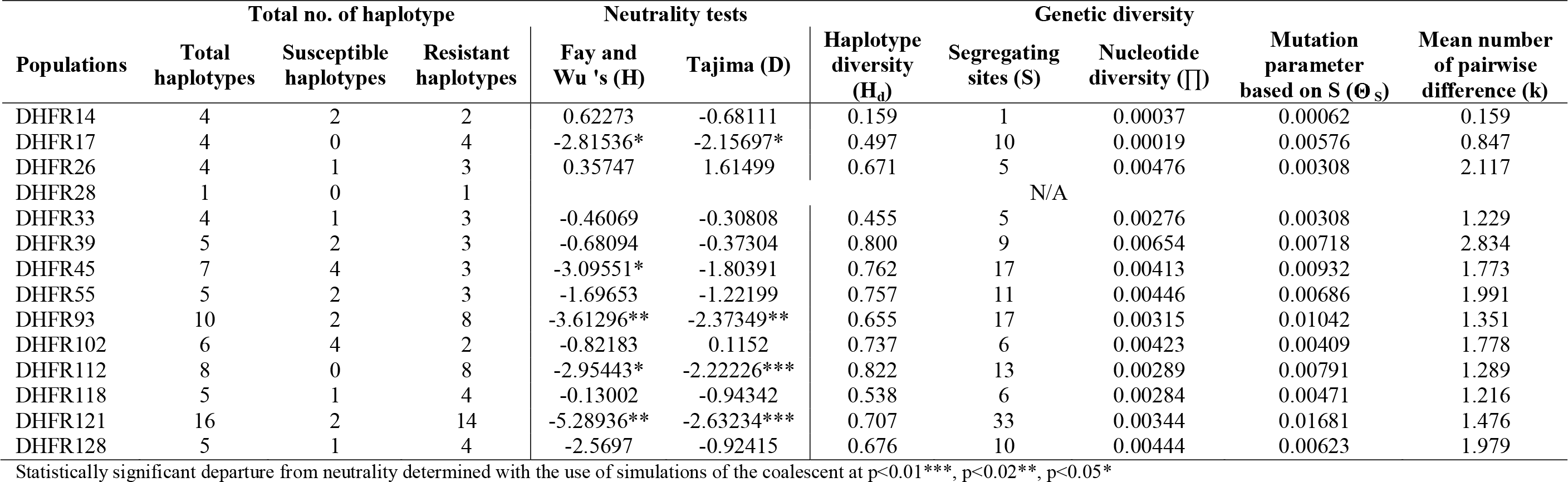
Total number of susceptible and resistant haplotype, neutrality analysis and genetic diversity of dihydrofolate reductase locus from fourteen *P. vivax* populations.

### 3.4 Distribution of P. vivax dihydrofolate reductase haplotypes

A total 42 unique haplotypes of the dihydrofolate reductase locus were identified among all 14 populations. Twenty-one haplotypes encoded S58R (AG<u>A</u>)/S117N (A<u>A</u>C) resistance mutations, 8 haplotypes encoded S117N (A<u>A</u>C) and one haplotype encoded either S58R (AG<u>A</u>) or I173L (<u>C</u>TT) resistance mutations (Figs 3 and 4). Based on the analysis of each population separately, three populations (DHFR93, DHFR112, DHFR121) contained maximum of 8 and 14 S58R (AG<u>A</u>)/S117N (A<u>A</u>C) and S117N (A<u>A</u>C) resistance haplotypes (Table 2, Supplementary Table S3). Three populations (DHFR17, DHFR118, DHFR128) contained maximum of 4 S58R (AG<u>A</u>)/S117N (A<u>A</u>C) and S117N (A<u>A</u>C) resistance haplotypes, 5 populations (DHFR26, DHFR33, DHFR39, DHFR45, DHFR55) contained maximum of 3 S58R (AG<u>A</u>)/S117N (A<u>A</u>C), S117N (A<u>A</u>C), S58R (AG<u>A</u>) and I173L (<u>C</u>TT) resistance haplotypes and 2 populations (DHFR14, DHFR102) contained maximum of 2 S58R (AG<u>A</u>)/S117N (A<u>A</u>C) and S117N (A<u>A</u>C) resistance haplotypes. A single I173L (<u>C</u>TT) resistance-conferring haplotype was present in the DHFR28 population (Table 2, Supplementary Table S3). Overall, the results show multiple or single resistance haplotypes in each population, demonstrating evidence of both hard and soft selective sweep patterns.

**Fig. 3.**
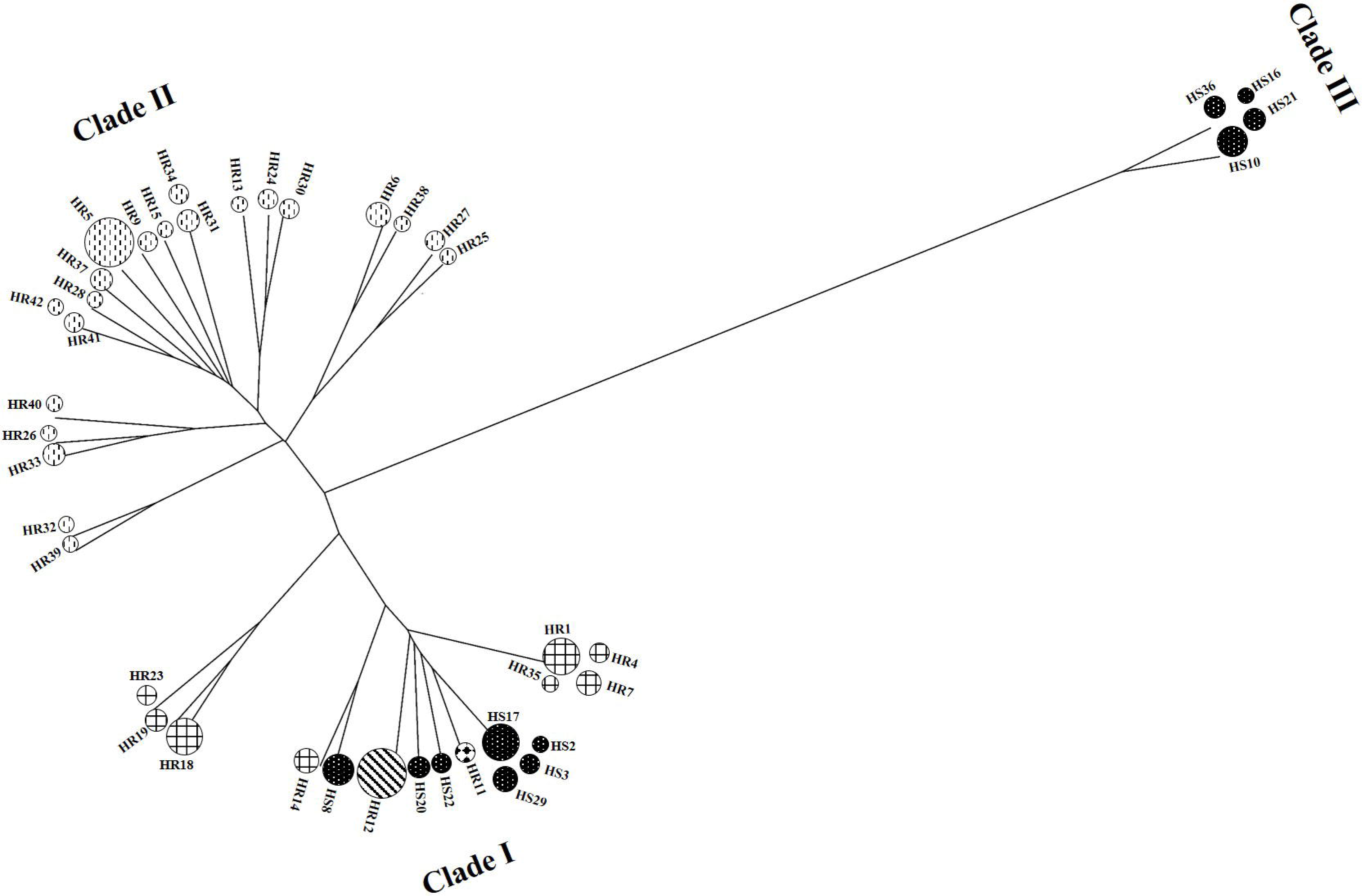
Split tree of the forty-two dihydrofolate reductase haplotypes was generated with the UPGMA method using the Jukes Cantor (JC+G) model of substitution in SplitsTrees4 software (Huson and Bryant, 2006). The circles represent the different haplotypes and the size of each circle is proportional to the number of sequences generated in that haplotypes from fourteen populations. The mutation carried each haplotype was shaded as follows: eleven susceptible haplotypes containing S58R(A<u>G</u>C)/ S117N(A<u>G</u>C)/ I173L (A<u>T</u>T) SNPs shown by black shading; eight P117N resistant haplotypes containing S117N(A<u>A</u>C)/ S58R(A<u>G</u>C)/ I173L (A<u>T</u>T) SNPs were hatched line shading; twenty one P58R/P117N double mutant resistant haplotypes containing S58R(AG<u>A</u>)/ S117N(A<u>A</u>C)/ I173L (A<u>T</u>T) SNPs shown by vertical dots shading; one P173L resistant haplotype containing I173L (<u>C</u>TT)/ S58R(A<u>G</u>C)/ S117N(A<u>G</u>C) SNPs shown by diagonal line shading and one S58R resistant haplotype containing S58R(A<u>G</u>A)/ S117N(A<u>G</u>C)/ I173L (A<u>T</u>T) SNPs have solid diamond shading.

**Fig. 4.**
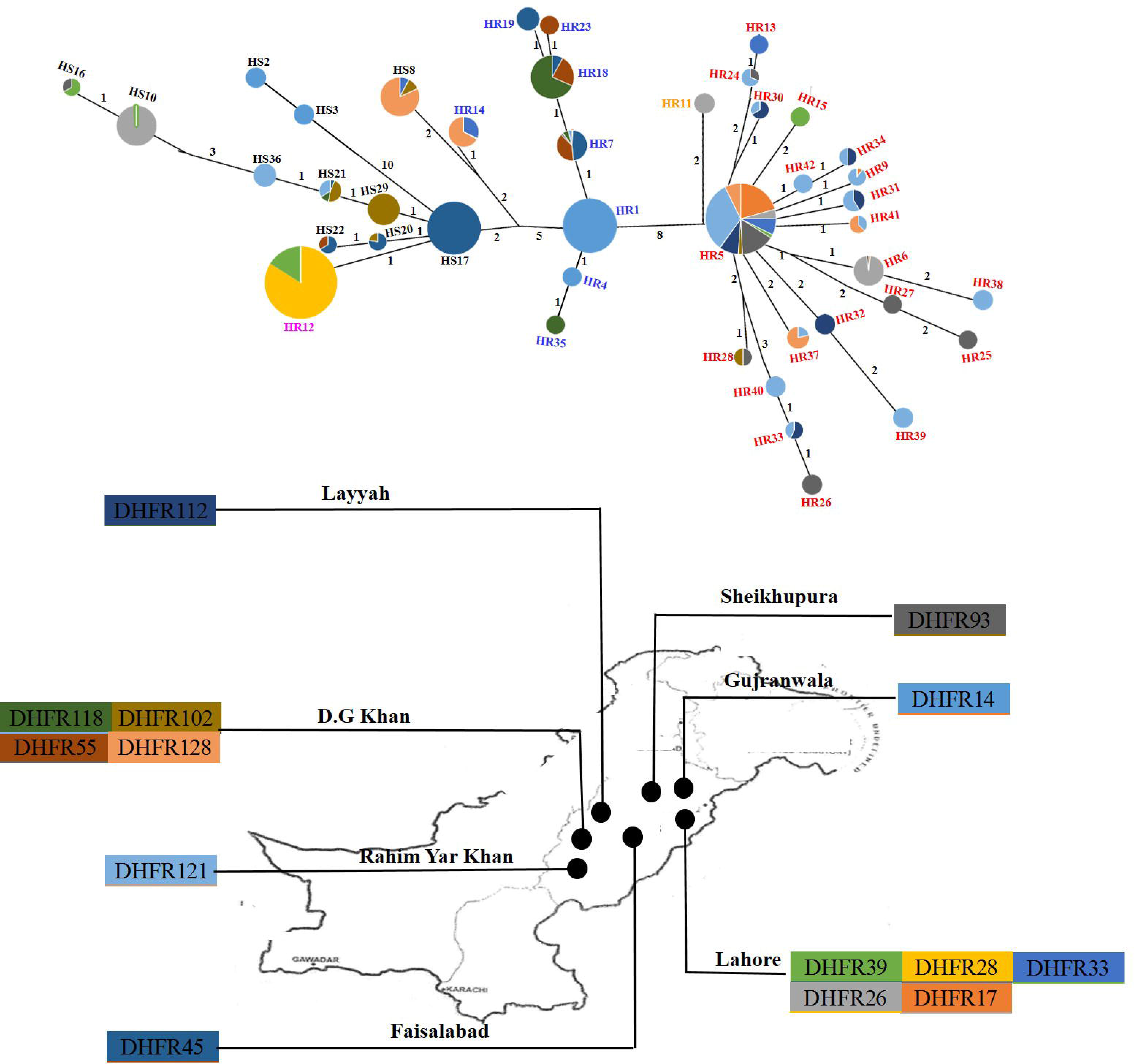
Network tree of the forty-two dihydrofolate reductase haplotypes was generated with the Neighbour Joining method in the Network 4.6.1 software (Fluxus Technology Ltd.). All unnecessary median vectors and links were removed with the star contractions. The size of each circle represents the haplotype was proportional to the number of sequences generated from different populations. The colors in the pie chart circles replicate the haplotype frequency and their distribution come from each of the fourteen populations as indicated on insert map. The number of mutations separating adjacent sequence nodes was indicated along connecting branches and the length of the lines connecting the haplotypes is proportional to the number of nucleotide changes. The mutation carried each haplotype was colored as follows: eleven susceptible haplotypes (HS) containing S58R(A<u>G</u>C)/ S117N(A<u>G</u>C)/ I173L (A<u>T</u>T) SNPs shown in black; eight P117N resistant haplotypes (HR) containing S117N(A<u>A</u>C)/ S58R(AGC)/ I173L (ATT) SNPs shown in blue; twenty one P58R/P117N double mutant resistant haplotypes (HR) containing S58R(AG<u>A</u>)/ S117N(A<u>A</u>C)/ I173L (A<u>T</u>T) SNPs shown in red; one P173L resistant haplotype (HR) containing I173L (C<u>T</u>T)/ S58R(A<u>G</u>C)/ S117N(AGC) SNPs shown in pink; one S58R resistant haplotype (HR) containing S58R(A<u>G</u>A)/ S117N(A<u>G</u>C)/ I173L (A<u>T</u>T) SNPs shown in yellow.

### 3.5 Phylogenetic relationships between pyrimethamine resistance associated SNPs in P. vivax populations

The split and network trees were produced to examine the phylogenetic relationship between forty-two dihydrofolate reductase haplotypes. The resistance haplotypes were distributed across the trees consistent with the emergence of resistance from different susceptible ancestral backgrounds (Fig. 3 and 4). When analysing the individual resistance mutations, the split tree of the S58R (AG<u>A</u>)/S117N (A<u>A</u>C) mutation reveals that twenty-one haplotypes (HR1, HR39, HR32, HR33, HR26, HR40, HR41, HR42, HR28, HR37, HR9, HR15, HR34, HR31, HR13, HR24, HR30, HR6, HR38, HR27, HR25) were located in a single distinct clade II (Fig. 3), indicating a single origin for this mutation (Supplementary Fig. S1). Susceptible haplotypes were not share with resistance haplotypes in clade II but arose separately in the closest clade III (Fig. 3). The network tree analysis showed that the 21 resistant haplotypes were present in eleven different populations collected from five major cities of Lahore, Sheikhupura, D.G. Khan, Layyah and R.Y. Khan (Fig. 4). Only one haplotype (HR5) was present at a high frequency in all 11 different populations from the five cites, showing its spread between the cities (Fig. 4).

For the S117N (A<u>A</u>C) mutation, the split tree analysis showed 8 resistant haplotypes (HR1, HR7, HR18, HR4, HR35, HR19, HR23, HR14) located in single clade I (Fig. 3), again indicating a single origin of this mutation (Supplementary Fig. S1). Each of these resistance haplotypes was closely related to one or more susceptible haplotypes. The network tree analysis showed that the 8 resistance associated haplotypes were present in 9 different populations collected from six major cities of Gujranwala, Lahore, Faisalabad, D.G. Khan, Sheikhupura and Layyah (Fig. 4). Three haplotypes (HR1, HR7, HR18) were present at a high frequency in the data set from all 8 different populations samples from six cites. The HR1 resistance haplotype was present only in a single population (DHFR14) collected from Gujranwala city, indicating that this mutation has not spread to any other city. The HR7 and HR18 haplotypes were present in three and six different populations respectively, sampled from the five cites excluding Gujranwala (Fig. 4), indicating the spread of this mutations between these cities (Fig. 4). The overall phylogenetic relationship supports the view that S58R (AG<u>A</u>)/S117N (A<u>A</u>C) and S117N (A<u>A</u>C) mutations arose at multiple times from a single origin and spread to the multiple different locations in the Punjab province of Pakistan through the flow of drug resistance alleles.

Interestingly, the I173L (<u>C</u>TT) resistance SNP was present on a single (HR12) haplotype located in clade I of the split tree (Fig. 3, Supplementary Fig. S1). This resistance haplotype was more closely related to one or more susceptible haplotypes and represented in two populations (DHFR28, DHFR39) from the city of Lahore (Fig. 4). The data showed that HR12 haplotype was present at a high frequency in both populations (Fig. 4). In conclusion, I173L (<u>C</u>TT) mutation was present on a single haplotype and the phylogenetic relationship suggests that it arose rarely and has not spread to different cities.

## 4. Discussion

The frequency with which pyrimethamine resistance emerges and the extent to which it spreads are the important considerations for its prevention and control. Our study used molecular genetic approaches to provide relevant insight to the emergence and spread of resistance after intensive positive selection pressure at the dihydrofolate reductase locus in the Punjab province of Pakistan. We chose this region because patients are treated in a sporadic manner often with generic pyrimethamine drugs of unknown quality.

### 4.1 *Pyrimethamine resistance mutations in* P. vivax *and their effect on positive selection pressure*

Pyrimethamine resistance has been investigated in a number of different geographical regions and strong evidence exists that F57L/I (TT<u>A</u>/<u>A</u>T<u>A</u>), S58R (AG<u>A</u>), T61M (A<u>T</u>G), S117N/T (A<u>A</u>C/A<u>C</u>C) and I173L/F (<u>C</u>TT/<u>T</u>TT) SNPs are responsible for resistance in *P. vivax* (Hawkins et al., 2007). Subsequent work has consistently shown that F57L/I (TT<u>A</u>/<u>A</u>T<u>A</u>), S58R (AG<u>A</u>), S117N/T (A<u>A</u>C/A<u>C</u>C) single mutations and S58R (AG<u>A</u>) /S117N/T (A<u>A</u>C/A<u>C</u>C) double mutations are widely distributed in different geographical regions of south Asia (Auliff et al., 2006; Brega et al., 2004; de Pecoulas et al., 1998; Hastings et al., 2005; Imwong et al., 2003; Kaur et al., 2006; Kuesap et al., 2011; Lu et al., 2012; Mint Lekweiry et al., 2012; Ranjitkar et al., 2011; Schunk et al., 2006). Previous studies in Pakistan have shown that the S58R (AG<u>A</u>)/S117N (A<u>A</u>C) double mutation and S117N (A<u>A</u>C) single mutation conferring pyrimethamine resistance were present in different cities of the Punjab, Sindh and KPK provinces (Khattak et al., 2013; Zakeri et al., 2011). In the present study, the S58R (AG<u>A</u>)/ S117N (A<u>A</u>C) double mutation was also present in 10/38 populations. The single mutations of S117N (A<u>A</u>C) and S58R (AG<u>A</u>) were present in 8/38 and 1/38 populations, respectively. In contrast, a few studies have shown the I173L (<u>C</u>TT) mutation in south Asia (Auliff et al., 2006; Brega et al., 2004), but there was no evidence for the presence of this mutation in Pakistan. In the present study, we have identified for the first time the I173L (<u>C</u>TT) mutation in 2/38 populations. This mutation has previously been reported in just two populations of *P. vivax* collected from patients in Malo Island, Vanuatu (Auliff et al., 2006) and in a traveller on return to France from Afghanistan (Brega et al., 2004), but has not been detected in other surveys from south Asia.

Differences in the frequency of pyrimethamine conferring mutations between geographical regions may be explained by variable drug doses, for example if the I173L (<u>C</u>TT) mutations are only selected at low doses of pyrimethamine, while S58R (AG<u>A</u>) /S117N/T (A<u>A</u>C/A<u>C</u>C) double mutations occur at higher doses. An alternative explanation may be the mutations confer a fitness cost or disadvantage. This suggests that fitness effects due to pyrimethamine resistance may vary among different parasite populations due to their genetic background. A third explanation is that difference may be due to genetic drift or bottlenecking effects, for example, if changes in the weather disrupt the life cycle of *P. vivax* in mosquito result in to the removing of less common mutation. A fourth explanation is may be due to movement of patients to new places with pyrimethamine susceptible *P. vivax* removing less common mutations [e.g. I173L (<u>C</u>TT)] at mating with susceptible parasites.

### 4.2 Nature of selective sweeps associated with the emergence of pyrimethamine resistance alleles in P. vivax

Several studies have assessed the selective sweep patterns in pyrimethamine resistance mutations of *P. falciparum* at the dihydrofolate reductase locus from different geographical regions. These emphasise a reduction in polymorphism around the dihydrofolate reductase locus with a single resistance haplotype indicative of a hard selective sweeps (McCollum et al., 2007; Nair et al., 2003; Pearce et al., 2005; Roper et al., 2003). A few studies have examined the presence of multiple haplotypes in Kenyan and Cameroon populations of *P. falciparum* indicative of soft selective sweeps (McCollum et al., 2008; McCollum et al., 2006). There is a single report for a selective advantage to parasites bearing resistance conferring mutations at the *P. vivax* dihydrofolate reductase, providing evidence for a high level of polymorphism around the locus with multiple resistance haplotypes indicative of a soft selective sweeps (Hawkins et al., 2008). However, there is currently little understanding of the nature of selective sweeps associated with the emergence of pyrimethamine resistance in *P. vivax*. In the present study, although both soft and hard selective sweeps were present, there is a predominance of hard sweeps (Fig.1). A single resistance haplotype at high frequency was detected in the DHFR28 population and 2 to 3 resistance haplotypes were present in eight populations (DHFR55, DHFR128, DHFR118, DHFR102, DHFR14, DHFR39, DHFR33, DHFR26), only one of these predominating at high frequency (Fig. 1). The selective sweeps on these nine individual populations were effectively hard with no evidence of the genetic footprint of selection. In contrast, there were 4 to 8 resistance haplotypes at high frequency in five populations (DHFR112, DHFR121, DHFR45, DHFR93, DHFR17) (Fig. 1). The selective sweep on these five individual populations was effectively soft and a genetic footprint of selection was also detected by significant departures from neutrality test (Table 1).

It is much easier to identify the emergence of resistance mutations with a single haplotype at high frequency, characteristic of a hard selective sweeps. Our results are consistent with the hypothesis that a single mutation with high frequency emerged in nine populations (Fig. 1). It can be more difficult to demonstrate the emergence of resistance mutations with multiple haplotypes at high frequency due to recurrent mutations after the onset of selection or pre-existing mutations before the onset of selection. Pre-exiting mutations are seen in multiple haplotypes after historical recombination, and if selected, lead to a high level of haplotype diversity, whereas recurrent mutations appear in populations after the onset of selection (Redman et al., 2015). If pre-existing mutations were the only source of emergence of resistance mutations, similar resistance haplotypes would be present in many populations, even if resistance alleles do not shared between those populations and were present prior the onset of selection (Chaudhry et al., 2016). This is not the case in the present study, where there is high level of gene flow between those populations, still DHFR45 and DHFR17 populations carried different resistance haplotypes (Fig. 1), providing strong evidence for the emergence of resistance by multiple independent recurrent mutations.

### 4.3 Role of human travel in the spread of pyrimethamine resistance mutations

There is large amount of human movement in the Punjab province of Pakistan, hence migration may play an important role in the spread of resistance mutations. However, the presence of same resistance haplotypes in two or three populations cannot itself be taken as conclusive evidence of the spread of resistance mutations (Redman et al., 2015). Nonetheless, the present study suggests that human migration between cities is an important factor in the spread of pyrimethamine resistance in Punjab. The S58R (AG<u>A</u>)/S117N (A<u>A</u>C) and S117N (A<u>A</u>C) mutations were present on multiple diverse resistance haplotypes and their phylogenetic relationship suggests that there is a single independent origin of these two mutations in the populations examined (Fig. 2). It is notable that the HR5 resistance haplotype of the S58R (AG<u>A</u>)/S117N (A<u>A</u>C) mutation predominates in eleven different populations sampled from five cities of Punjab and the other two resistance haplotypes (HR18 and HR 7) of S117N (A<u>A</u>C) mutation were present in three and six different populations respectively, sampled from five cities of Punjab (Fig. 3). It is likely that human travelling has contributed to the spread of these resistance haplotypes in the Punjab province from a single origin. Interestingly in the case of the I173L (<u>C</u>TT) mutation, our analysis suggests that it arose rarely from a single origin (Fig. 2). The mutation was present on a single haplotype in two *P. vivax* populations within the city of Lahore (Fig. 3), supporting the hypothesis that it will spread in future, if control measures are not implemented.

## 5. Conclusion

Understanding of the emergence and the spread of pyrimethamine resistance mutations in *P. vivax* is relevant to the implementation of sustainable parasite control strategies. Our models suggests that once the resistance has been established, gene flow plays a key role in its spread. Therefore, a new drug combination should be introduced in the country, could potentially pose the spread of existing drug resistance alleles that have previously intensive positive selection pressure. Other priority for the management planning is the sensible use of drug, movement of infected patient, vector control strategies. These recommendations will be disseminated using appropriate educational methods and tools targeted to different stakeholder.

## Acknowledgements

We are grateful for the contributions of Sana Amir and Saqib Shahzad (Chugtai Laboratories Punjab Pakistan) for providing the opportunity to collect samples. We are grateful for the assistance and guidance of Dr. AR Shakoori (University of Central Punjab Pakistan). The study was financially supported by the University of Central Punjab Pakistan.

## Supplementary Figure Legends

**Supplementary Fig. S1.** Split tree of forty-two dihydrofolate reductase haplotypes sequenced from fourteen *P. vivax* populations. The tree was obtained by UPGMA analysis using the JukesCantor (JK+G+I). model of substitution (Drummond AJ, 2012). Branches with bootstrap support values above 50% (1000 replications). The phylogeny is rooted with a dihydrofolate reductase sequence of *Plasmodium falciparum*. The mutation carried each haplotype was shaded as follows: eleven susceptible haplotypes containing S58R(A<u>G</u>C)/ S117N(A<u>G</u>C)/ I173L (A<u>T</u>T) SNPs shown by black shading; eight P117N resistant haplotypes containing S117N(A<u>A</u>C)/ S58R(A<u>G</u>C)/ I173L (A<u>T</u>T) SNPs were hatched line shading; twenty one P58R/P117N double mutant resistant haplotypes containing S58R(AG<u>A</u>)/ S117N(A<u>A</u>C)/ I173L (A<u>T</u>T) SNPs shown by vertical dots shading; one P173L resistant haplotype containing I173L (<u>C</u>TT)/ S58R(A<u>G</u>C)/ S117N(A<u>G</u>C) SNPs shown by diagonal line shading and one S58R resistant haplotype containing S58R(A<u>G</u>A)/ S117N(A<u>G</u>C)/ I173L (A<u>T</u>T) SNPs have solid diamond shading.

## References

Alam, M.T., Agarwal, R., Sharma, Y.D., 2007. Extensive heterozygosity at four microsatellite loci flanking Plasmodium vivax dihydrofolate reductase gene. Mol Biochem Parasitol 153, 178–185.

Asif, S.A., 2008. Departmental audit of malaria control programme 2001-2005 north west frontier province (NWFP). J Ayub Med Coll Abbottabad 20, 98–102.

Auliff, A., Wilson, D.W., Russell, B., Gao, Q., Chen, N., Anh le, N., Maguire, J., Bell, D., O’Neil, M.T., Cheng, Q., 2006. Amino acid mutations in Plasmodium vivax DHFR and DHPS from several geographical regions and susceptibility to antifolate drugs. Am J Trop Med Hyg 75, 617–621.

Brega, S., de Monbrison, F., Severini, C., Udomsangpetch, R., Sutanto, I., Ruckert, P., Peyron, F., Picot, S., 2004. Real-time PCR for dihydrofolate reductase gene single-nucleotide polymorphisms in Plasmodium vivax isolates. Antimicrob Agents Chemother 48, 2581–2587.

Chaudhry, U., 2015. Molecular Analysis of Benzimidazole Resistance in Haemonchus contortus and Haemonchus placei. PhD Thesis

Chaudhry, U., E. M. Redman, Muthusamy Raman, Gilleard., J.S., 2015. Genetic evidence for the spread of a benzimidazole resistance mutation across southern India from a single origin in the parasitic nematode Haemonchus contortus International Journal for Parasitology IJPARA-S-15-00127-2.

Chaudhry, U., Redman, E.M., Ashraf, K., Shabbir, M.Z., Rashid, M.I., Ashraf, S., Gilleard, J.S., 2016. Microsatellite marker analysis of Haemonchus contortus populations from Pakistan suggests that frequent benzimidazole drug treatment does not result in a reduction of overall genetic diversity. Parasites & Vectors 9, 349.

Conway, D.J., 2007. Molecular epidemiology of malaria. Clin Microbiol Rev 20, 188–204.

de Pecoulas, P.E., Tahar, R., Ouatas, T., Mazabraud, A., Basco, L.K., 1998. Sequence variations in the Plasmodium vivax dihydrofolate reductase-thymidylate synthase gene and their relationship with pyrimethamine resistance. Mol Biochem Parasitol 92, 265–273.

Drummond AJ, A.B., Buxton S, Cheung M, Cooper A, Duran C, Field M, Heled J, Kearse M, Markowitz S, Moir R, Stones-Havas S, Sturrock S, Thierer T, Wilson A 2012. Geneious v5.6.

Fay, J.C., Wu, C.I., 2000. Hitchhiking under positive Darwinian selection. Genetics 155, 1405–1413.

Hastings, M.D., Maguire, J.D., Bangs, M.J., Zimmerman, P.A., Reeder, J.C., Baird, J.K., Sibley, C.H., 2005. Novel Plasmodium vivax dhfr alleles from the Indonesian Archipelago and Papua New Guinea: association with pyrimethamine resistance determined by a Saccharomyces cerevisiae expression system. Antimicrob Agents Chemother 49, 733–740.

Hastings, M.D., Porter, K.M., Maguire, J.D., Susanti, I., Kania, W., Bangs, M.J., Sibley, C.H., Baird, J.K., 2004. Dihydrofolate reductase mutations in Plasmodium vivax from Indonesia and therapeutic response to sulfadoxine plus pyrimethamine. J Infect Dis 189, 744–750.

Hastings, M.D., Sibley, C.H., 2002. Pyrimethamine and WR99210 exert opposing selection on dihydrofolate reductase from Plasmodium vivax. Proc Natl Acad Sci U S A 99, 13137–13141.

Hawkins, V.N., Auliff, A., Prajapati, S.K., Rungsihirunrat, K., Hapuarachchi, H.C., Maestre, A., O’Neil, M.T., Cheng, Q., Joshi, H., Na-Bangchang, K., Sibley, C.H., 2008. Multiple origins of resistance-conferring mutations in Plasmodium vivax dihydrofolate reductase. Malaria Journal 7, 72.

Hawkins, V.N., Joshi, H., Rungsihirunrat, K., Na-Bangchang, K., Sibley, C.H., 2007. Antifolates can have a role in the treatment of Plasmodium vivax. Trends Parasitol 23, 213–222.

Huang, F., Zhou, S., Zhang, S., Li, W., Zhang, H., 2011. Monitoring resistance of Plasmdium vivax: Point mutations in dihydrofolate reductase gene in isolates from Central China. Parasites & Vectors 4, 80–80.

Huson, D.H., Bryant, D., 2006. Application of phylogenetic networks in evolutionary studies. Molecular biology and evolution 23, 254–267.

Imwong, M., Pukrittayakamee, S., Renia, L., Letourneur, F., Charlieu, J.P., Leartsakulpanich, U., Looareesuwan, S., White, N.J., Snounou, G., 2003. Novel point mutations in the dihydrofolate reductase gene of Plasmodium vivax: evidence for sequential selection by drug pressure. Antimicrob Agents Chemother 47, 1514–1521.

Kaur, S., Prajapati, S.K., Kalyanaraman, K., Mohmmed, A., Joshi, H., Chauhan, V.S., 2006. Plasmodium vivax dihydrofolate reductase point mutations from the Indian subcontinent. Acta Trop 97, 174–180.

Khattak, A.A., Venkatesan, M., Khatoon, L., Ouattara, A., Kenefic, L.J., Nadeem, M.F., Nighat, F., Malik, S.A., Plowe, C.V., 2013. Prevalence and patterns of antifolate and chloroquine drug resistance markers in Plasmodium vivax across Pakistan. Malar J 12, 1475–2875.

Kuesap, J., Rungsrihirunrat, K., Thongdee, P., Ruangweerayut, R., Na-Bangchang, K., 2011. Change in mutation patterns of Plasmodium vivax dihydrofolate reductase (Pvdhfr) and dihydropteroate synthase (Pvdhps) in P. vivax isolates from malaria endemic areas of Thailand. Mem Inst Oswaldo Cruz 1, 130–133.

Lee, W.J., Kim, H.H., Choi, Y.K., Choi, K.M., Kim, M.A., Kim, J.Y., Sattabongkot, J., Sohn, Y., Kim, H., Lee, J.K., Park, H.S., Lee, H.W., 2010. Analysis of the dihydrofolate reductase-thymidylate synthase gene sequences in Plasmodium vivax field isolates that failed chloroquine treatment. Malar J 9, 1475–2875.

Librado, P., Rozas, J., 2009. DnaSP v5: a software for comprehensive analysis of DNA polymorphism data. Bioinformatics 25, 1451–1452.

Lu, F., Wang, B., Cao, J., Sattabongkot, J., Zhou, H., Zhu, G., Kim, K., Gao, Q., Han, E.T., 2012. Prevalence of drug resistance-associated gene mutations in Plasmodium vivax in Central China. Korean J Parasitol 50, 379–384.

McCollum, A.M., Basco, L.K., Tahar, R., Udhayakumar, V., Escalante, A.A., 2008. Hitchhiking and Selective Sweeps of <em>Plasmodium falciparum </em> Sulfadoxine and Pyrimethamine Resistance Alleles in a Population from Central Africa. Antimicrobial Agents and Chemotherapy 52, 4089.

McCollum, A.M., Mueller, K., Villegas, L., Udhayakumar, V., Escalante, A.A., 2007. Common origin and fixation of Plasmodium falciparum dhfr and dhps mutations associated with sulfadoxine-pyrimethamine resistance in a low-transmission area in South America. Antimicrob Agents Chemother 51, 2085–2091.

McCollum, A.M., Poe, A.C., Hamel, M., Huber, C., Zhou, Z., Shi, Y.P., Ouma, P., Vulule, J., Bloland, P., Slutsker, L., Barnwell, J.W., Udhayakumar, V., Escalante, A.A., 2006. Antifolate resistance in Plasmodium falciparum: multiple origins and identification of novel dhfr alleles. J Infect Dis 194, 189–197.

Mint Lekweiry, K., Ould Mohamed Salem Boukhary, A., Gaillard, T., Wurtz, N., Bogreau, H., Hafid, J.E., Trape, J.F., Bouchiba, H., Ould Ahmedou Salem, M.S., Pradines, B., Rogier, C., Basco, L.K., Briolant, S., 2012. Molecular surveillance of drug-resistant Plasmodium vivax using pvdhfr, pvdhps and pvmdr1 markers in Nouakchott, Mauritania. J Antimicrob Chemother 67, 367–374.

Nair, S., Williams, J.T., Brockman, A., Paiphun, L., Mayxay, M., Newton, P.N., Guthmann, J.P., Smithuis, F.M., Hien, T.T., White, N.J., Nosten, F., Anderson, T.J., 2003. A selective sweep driven by pyrimethamine treatment in southeast asian malaria parasites. Molecular biology and evolution 20, 1526–1536.

Pearce, R., Malisa, A., Kachur, S.P., Barnes, K., Sharp, B., Roper, C., 2005. Reduced variation around drug-resistant dhfr alleles in African Plasmodium falciparum. Molecular biology and evolution 22, 1834–1844.

Petersen, I., Eastman, R., Lanzer, M., 2011. Drug-resistant malaria: molecular mechanisms and implications for public health. FEBS Lett 585, 1551–1562.

Posada, D., 2008. JModelTest: phylogenetic model averaging. Mol. Biol. Evol. 25, 1253–1256.

Ranjitkar, S., Schousboe, M.L., Thomsen, T.T., Adhikari, M., Kapel, C.M., Bygbjerg, I.C., Alifrangis, M., 2011. Prevalence of molecular markers of anti-malarial drug resistance in Plasmodium vivax and Plasmodium falciparum in two districts of Nepal. Malar J 10, 75.

Redman, E., Whitelaw, F., Tait, A., Burgess, C., Bartley, Y., Skuce, P., Jackson, F., Gilleard, J., 2015. The emergence of resistance to the benzimidazole anthlemintics in parasitic nematodes of livestock is characterised by multiple independent hard and soft selective sweeps. PLoS Negl Trop Dis. 6;9, :e0003494. doi:.

Roper, C., Pearce, R., Bredenkamp, B., Gumede, J., Drakeley, C., Mosha, F., Chandramohan, D., Sharp, B., 2003. Antifolate antimalarial resistance in southeast Africa: a population-based analysis. Lancet (London, England) 361, 1174–1181.

Schunk, M., Kumma, W.P., Miranda, I.B., Osman, M.E., Roewer, S., Alano, A., Loscher, T., Bienzle, U., Mockenhaupt, F.P., 2006. High prevalence of drug-resistance mutations in Plasmodium falciparum and Plasmodium vivax in southern Ethiopia. Malar J 5, 1475–2875.

Shaukat, M. Azmat Ullah Khan, Qasim Ali, Mushtaq A. Saleem, Neil D. Sargison, Timothy Connelley, Chaudhry, U., 2018. Highly sensitive illumina MiSeq deep amplicon sequencing technology for the identification of Plasmodium vivax and Plasmodium falciparum Manuscript In-preparation.

Tajima, F., 1989. Statistical method for testing the neutral mutation hypothesis by DNA polymorphism. Genetics 123, 585–595.

Tamura, K., Peterson, D., Peterson, N., Stecher, G., Nei, M., Kumar, S., 2011. MEGA5:Molecular Evolutionary Genetics Analysis using Maximum Likelihood, Evolutionary Distance, and Maximum Parsimony Methods. Mol. Biol. Evol. 28 (10), 2731–2739.

Zakeri, S., Afsharpad, M., Ghasemi, F., Raeisi, A., Kakar, Q., Atta, H., Djadid, N.D., 2011. Plasmodium vivax: prevalence of mutations associated with sulfadoxine-pyrimethamine resistance in Plasmodium vivax clinical isolates from Pakistan. Exp Parasitol 127, 167–172.

